# Bidirectional effects of anxiety and anorexia nervosa: A Mendelian randomization study

**DOI:** 10.1101/451500

**Authors:** E Caitlin Lloyd, Hannah Sallis, Bas Verplanken, Anne M Haase, Marcus R Munafò

## Abstract

**Objectives:** To assess bidirectional effects of anxiety and anorexia nervosa (AN) phenotypes. **Design** Two-sample Mendelian randomization.

**Setting:** Genome-wide association study (GWAS) summary statistics from the Psychiatric Genomics Consortium (PGC), analysis of the UK Biobank sample, and Anxiety Neuro Genetics Study (ANGST) consortium.

**Participants:** European descent participants from the PGC (n = 14,477), UK Biobank (n = 348,219), and ANGST consortium (n = 17,310, and n = 18,186).

**Main outcome measures:** AN diagnosis, worry, anxiety disorder pathology (case-control and quantitative phenotypes).

**Results:** We found evidence of a moderate genetic correlation between worry and AN (Rg = 0.36, SE = 0.05, p < 0.001), and the Mendelian randomization analysis supported a causal influence of worry on AN (OR = 2.14, 95% CI: 1.18 to 3.90, p = 0.01). There was no clear evidence for a causal effect of AN on worry in this study (B = −0.01, 95% CI: −0.03 to 0.02, p = 0.55). There was no robust evidence for a causal influence of anxiety disorders on AN (for case-control anxiety disorder phenotype: OR = 1.02, 95% CI: 0.69, 1.50, p = 0.922; for quantitative anxiety disorder phenotype: OR = 4.26, 95% CI: 0.49, 36.69, p = 0.187). There was no robust evidence for a causal effect of AN on anxiety disorders (for case control anxiety disorder phenotype: OR = 1.00, 95% CI: 0.72, 1.38, p = 0.981; for quantitative anxiety disorder phenotype: B = 0.01, 95% CI: −0.06, 0.6=09, p = 0,761). AN and anxiety disorder phenotypes were not genetically correlated (for case-control anxiety disorder phenotype: Rg = 0.10, se = 0.17, p = .56; for quantitative anxiety disorder phenotype: Rg = 0.12, SE = 0.17, p = 0.47).

**Conclusions:** Findings support a role for worry in AN development, highlighting a potential target of future AN prevention efforts. Mechanisms underlying the association should be a focus of future investigation. The relatively small sample sizes of anxiety disorder and AN GWASs may have limited power to detect causal effects; these associations should be studied further.

## Introduction

Anorexia nervosa (AN) is a serious eating disorder that is characterised by persistent restriction of caloric intake and fear of weight-gain in the context of a low body weight (1). AN has a lifetime prevalence rate of approximately 1 to 4% (2, 3), a range of lasting physical health complications (4), and the highest mortality rate of any psychiatric disorder (5). No single treatment or set of treatments has been found to be consistently successful, with AN recovery rates following treatment below 50% (6).

The scope for targeting putative mechanisms of AN is currently limited. Despite substantial development in the study of AN, with investigations focusing on a range of possible mechanisms (e.g. genetic, neural, psychological and personality factors), the aetiology of the disorder remains largely unknown (7). A number of models of illness have proposed a causal role of anxiety that does not surround eating and weight-gain (i.e., anxiety that is not explained by a diagnosis of AN) in the development of AN (8-11). Empirical evidence has provided some support for such models. Trait anxiety, a proneness to experiencing anxiety generally, is reported to be higher in individuals with AN as compared to healthy controls (12-14), and anxiety disorder prevalence is elevated in AN populations, as compared to the general population (15, 16). Importantly, retrospective studies report both anxious temperament and anxiety disorder pathology to precede the onset of AN (17-20), although findings from prospective studies are mixed (21, 22).

Although current evidence generally is consistent with a causal effect of anxiety on AN, the reported associations are at risk of confounding by unmeasured, or inadequately measured, factors. Demographic characteristics or other psychiatric comorbidities may increase risk for both anxiety and AN, serving to induce a correlation between the two, in the absence of a causal relationship. Reverse causation is also a possibility, with observed associations being driven by AN influencing anxiety, rather than the other way around. The association between malnutrition and anxiety in AN is currently unclear (23). However, nutrition affects various hormonal and neurotransmitter systems implicated in anxiety, changes to which have been observed in AN (24-27), and dietary restriction results in psychological and emotional changes in populations without AN (28, 29). A recent prospective study also found AN to increase the likelihood of a later anxiety disorder diagnosis (30). The biases that studies using traditional epidemiologic methods are subject to (e.g. confounding and reverse causation) mean that it is difficult to draw strong conclusions concerning the causal role of anxiety in AN using the existing evidence. However, being able to make confident inferences would better inform models of illness and the subsequent development of novel prevention and treatment interventions.

Mendelian randomization (MR) is an epidemiological approach that minimises bias affecting traditional observational epidemiology (31-33). The method uses genetic variants that are associated with the exposure of interest (in this case, anxiety) as instruments for examining the association between exposure and outcome (Figure 1). The association of the genetic variant with the outcome is analysed, under the assumption that the effect of the genetic variant is fully mediated by the exposure. This assumption is violated when horizontal pleiotropy occurs, that is, when the genetic variant is associated with other traits that also affect the outcome. Methods robust to this form of pleiotropy, and violations of other MR assumptions, have been developed. Consistency between estimates using these different methods can strengthen conclusions from MR studies (34).

**Figure 1:**
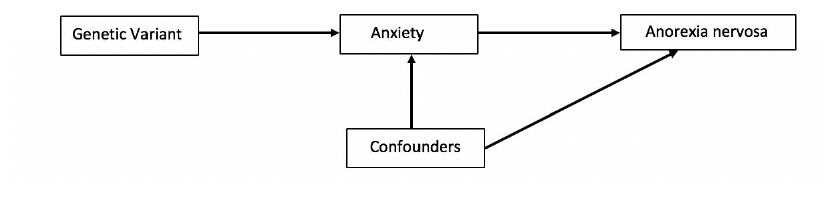
Diagram of a Mendelian randomization analysis

Mendel’s laws of segregation and independent assortment describe the random allocation of alleles during gamete formation. An individual’s genotype is the result of two such randomised transmissions: one maternal, and one paternal. The result is that genetic variants associated with the exposure of interest are generally not associated with traits that may confound the exposure-outcome association in traditional observational studies (35). Associations of a given genetic variant with the outcome of interest cannot be explained by reverse causation either, since the genotype one is born with is not altered by a disease outcome.

Where genetic variants are robustly associated with an exposure of interest, individuals with the risk increasing form of the variant will on average have greater levels of the exposure. However, groups will not differ with regard to confounding factors. MR is not concerned with making conclusions about the genetic underpinnings of an outcome, but rather with establishing an unbiased estimate of the effect of an exposure on an outcome, using genetic variants as proxy variables to achieve this (32). In a two-sample MR analysis an estimate of the association of the genetic variant with both the exposure and outcome is obtained. Gene-exposure associations are estimated in a different sample to the gene-outcome associations, meaning summary statistics from different genome-wide association studies (GWAS) may be used to complete the analysis. This approach will yield valid estimates providing the two samples are from the same underlying population (36).

A bidirectional MR analysis of worry and AN has been completed previously (37). Worry is defined as a negatively valanced and uncontrollable thought process, intended to resolve an issue that has at least one possible negative outcome (38). Worry is conceptualised as the cognitive component of anxiety (39), correlates highly with trait anxiety (40), is present in a number of anxiety disorders and is a core symptom of Generalised Anxiety Disorder (1). The existing MR study found no evidence of a causal association between worry and AN in either direction, although the two were genetically correlated (37). The AN GWAS included a relatively small number of cases however, which will have resulted in low sensitivity to detect causal effects of worry on AN (41), and vice versa (42), in the MR analysis.

Here we used summary data from the largest GWAS of AN completed to date (43) to investigate causal effects between anxiety and AN using genetic correlation and bidirectional two sample MR approaches. We extend previous investigations by considering the association of anxiety disorder phenotypes, in addition to worry, with AN. Findings from observational studies suggest the existence of causal influences in both directions, supporting the notion of a cycle in which anxiety is relieved by dietary restriction, but then elevated beyond initial levels to prompt further starvation (9, 44). We therefore hypothesised that we would observe bidirectional effects.

## Method

### Data sources

Details of the GWAS data used in the current study are provided in Table 1. The worry phenotype was quantitative, and measured by items comprising the worry dimension of the Eysenck personality questionnaire short-form neuroticism subscale (45, 46), that was administered to participants of the UK Biobank study. Binary responses (yes/no) to the questions ‘Are you a worrier?’, ‘Do you suffer from nerves?’, ‘Would you call yourself a nervous person?’ and ‘Would you call yourself tense or highly strung’, were summed to create a total score out of four, with higher scores indicating more severe worry. Only individuals who provided valid responses to all items were included in the GWAS. The cluster of worry items are reported to display a distinct genetic signal, in comparison to other clusters of the neuroticism subscale (37).

The anxiety disorder case control phenotype reflects the presence of five core anxiety disorder pathologies (GAD, PD, social phobia, agoraphobia, specific phobia). Only individuals with threshold pathologies or no pathology were included to increase genetic signal. The quantitative anxiety disorder phenotype indicates liability for a common dimension of anxiety, and was developed from modelling covariation across the same five disorders (47). The AN phenotype was binary, and indicated lifetime AN, or eating disorder not otherwise specified AN subtype, diagnosis (43)

**Table 1:** GWAS Study Characteristics **Phenotype Study Resource Sample size Population Data Source**

| Phenotype | Study | Resource | Sample size | Population | Data Source |
| --- | --- | --- | --- | --- | --- |
| Worry | Nagel et al. 2018 (37) | UK Biobank | 348,219 | European | https://ctg.cncr.nl/software/summary_statistics |
| Anxiety Disorder (Case Control) | Ottawa et al. 2016 (47) | ANGST | 5712 cases<br>11598 controls | European | https://www.med.unc.edu/pgc/results-and-downloads |
| Anxiety Disorder (Quantitative) | Ottawa et al. 2016 (47) | ANGST | 18186 | European | https://www.med.unc.edu/pgc/results-and-downloads |
| Anorexia Nervosa | Duncan et al. 2017 (43) | PGC | 3495 Cases<br>10982 Controls | European | https://www.med.unc.edu/pgc/results-and-downloads |

ANGST = Anxiety NeuroGenetics Study Consortium; PGC = Psychiatric Genomics Consortium

### Genetic Instrument selection

Genetic instruments for each exposure of interest were identified from relevant GWAS statistics (Table 1). We initially used a significance threshold of 5 x 10^-8^ to select single nucleotide polymorphisms (SNPs) for use as instruments, to ensure robust associations between SNPs and each exposure (48). SNPs were clumped to ensure independence using a threshold of LD r^2=^0.001, and a distance of 10000kb. Where instruments comprised a single SNP following clumping, we ran an additional sensitivity analysis using a significance threshold of 5 x 10^-6^ for instrument identification.

If palindromic SNPs were indicated for eligible instruments, proxy variants were identified with the package proxysnps (49), using an R^2^ threshold of > 0.8, and LD scores from the European 1000 Genomes data. Where instrumental SNPs were missing from the outcome GWAS, proxy variants were identified using the same approach, and replaced original instruments for estimation of instrument-outcome associations where possible. Proxy variant details are provided in Table 1 of the Supplementary Material. The inclusion of proxies did not affect the independence of instrumental SNPs.

There were 60 SNPs associated with the worry exposure, 57 of which (or proxies) were available in the AN GWAS. Anxiety disorder and AN instruments included one independent SNP following clumping. When the SNP-exposure threshold was reduced, seven SNPs were associated with the anxiety disorder case control phenotype, and nine with the quantitative phenotype. The weaker AN instrument contained 16 independent SNPs; eleven were available in the worry GWAS, while eight were available in the anxiety disorder GWAS.

### Statistical Analyses

GWAS summary statistics were downloaded from consortium/study websites (Table 1) and converted into the format required for statistical analyses.

### Genetic Correlation Analyses

To estimate the genetic correlation between anxiety and AN phenotypes cross-trait linkage disequilibrium score regression (50) was implemented, using the ldsc command line tool (https://github.com/bulik/ldsc) and LD scores computed from the 1000 Genomes European data (https://data.broadinstitute.org/alkesgroup/LDSCORE/).

### Mendelian Randomization Analyses

Bidirectional MR analyses were implemented in R (51) using code available in the TwoSampleMR package of the analytical platform MR base (52), and local data. For single SNP instruments the Wald Ratio method, or the ratio of coefficients method (53), was used to estimate the causal effect. Where multiple SNPs were identified as eligible instruments, Wald ratio estimates for the different SNPs were combined in an inverse variance weighted (IVW) analysis (54). Cochrane’s Q statistic was calculated to assess the heterogeneity of estimates combined in the IVW analysis. Since the Q statistic is heavily affected by sample size, I^2^ and associated confidence intervals were also calculated, using formulae derived from the meta-analysis literature (55). ‘Leave one out’ analyses were completed when heterogeneity was detected: the IVW analysis was completed leaving out one SNP each time, and estimates plotted.

We completed three sensitivity analyses that are robust to horizontal pleiotropy, to evaluate the validity of IVW estimates. MR Egger regression (56) was used to estimate pleiotropic effects present in the IVW analysis, and provide a pleiotropy-corrected estimate of the causal effect. Rucker’s Q indicates heterogeneity around the Egger estimate (57), and was deducted from Cochrane’s Q; a large positive value, combined with evidence of pleiotropy, suggests the MR Egger model is a better fit to the data than the IVW model (57). Weighted median (58) and weighted mode (59) analyses, which provide consistent causal estimates when a proportion of genetic instruments are invalid, were also completed. For an overview of MR methods, see (60).

Where the MR analysis indicated a causal effect, we conducted a MR Steiger sensitivity analysis to evaluate whether the inferred direction of causal influence was correct. This estimates the variance explained in exposure and outcome for each variant, testing whether associations between genetic instruments and the exposure are stronger than corresponding associations between genetic instruments and the outcome. Where this is the case a direction of effect from exposure to outcome is supported (61). The MR analysis was replicated using the subsample of variants that showed stronger associations with the exposure as compared to the outcome.

### Estimate interpretation

The causal estimate reflects the change in outcome resulting from a unit change in exposure, and estimates for binary outcomes are exponentiated to reflect the increase in odds of an outcome per unit change in exposure. When the exposure is binary, estimates denote the change in outcome, or odds of outcome, per log-odds increase in the exposure.

## Results

### Genetic Correlation Analyses

Figure 2 displays the full results of the genetic correlation analyses. We found evidence that AN was genetically correlated with the worry phenotype: *Rg* = 0.36, SE = 0.05, *p* < 0.001. There was no strong evidence of a genetic association between AN and either anxiety disorder exposure.

**Figure 2:**
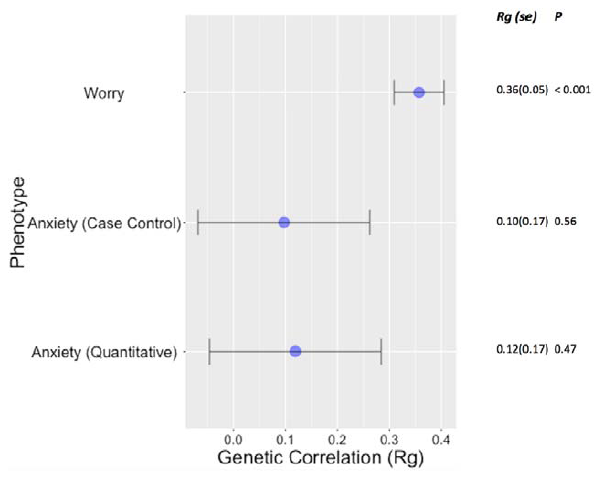
Genetic correlations between anxiety phenotypes and AN

### Mendelian Randomization Analyses

Bidirectional causal effects between worry/anxiety and AN phenotypes were assessed. Findings are summarised below.

### Causal influence of worry/anxiety disorders on AN

The IVW estimate indicated that worry increased the likelihood of AN diagnosis (OR = 2.14, 95% CI: 1.18, 3.90, p = 0.013). The weighted median estimate was consistent with this finding (OR = 2.49, 95% CI: 1.15, 5.41, p = 0.021), and the weighted mode estimate provided weak evidence for a positive association. The MR Egger estimate was not consistent with IVW, weighted median and weighted mode estimates, and confidence intervals around the estimate were very wide (Figure 3). Wald ratio estimates for each SNP are available in Figure 1 of Supplementary Material.

Outcomes of the MR Steiger investigation indicated that 37 of 57 variants showed stronger associations with the exposure as compared to the outcome (Supplementary Material, Table 2). Point estimates of the MR analysis using only these variants were consistent with those of the analysis including all 57 genetic instruments (i.e. supported worry increasing risk for AN), however the former were relatively imprecise (Supplementary Material, Figure 2).

There was no evidence for a causal influence of anxiety disorder pathology on AN in the single SNP analyses (Figure 3). Findings from sensitivity analyses that used multiple independent SNPs (less strongly associated with the anxiety disorder exposure) were consistent with those of single SNP analyses (Supplementary Material, Figures 3 and 4).

**Figure 3:**
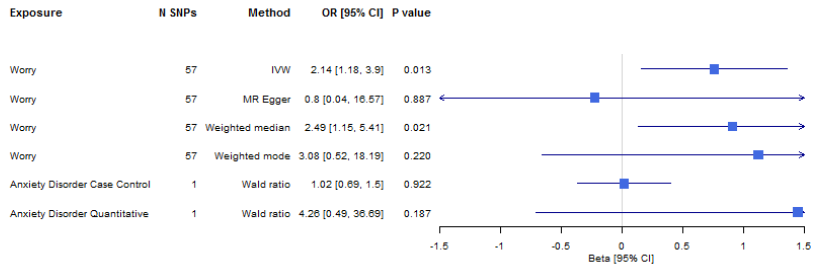
Mendelian randomization analysis to estimate causal influence of anxiety phenotypes on AN

### Causal influence of AN on worry/anxiety disorders

There was no strong evidence for a causal influence of AN on the worry phenotype, or either anxiety disorder phenotype, using the single SNP instrument (rs4622308) that was significant at the genome-wide level (Figure 4). Inferences from analyses using multiple SNP instruments did not qualitatively differ (Supplementary Material, Figures 5 - 7), with effect estimates remaining close to the null.

**Figure 4:**
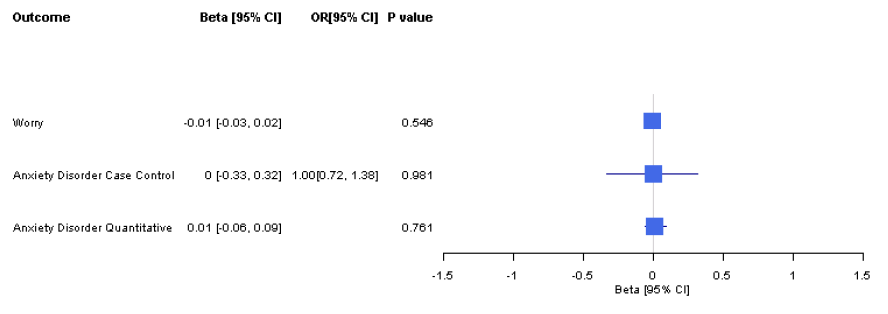
Mendelian randomization analysis to estimate causal influence of AN on anxiety phenotypes

### Pleiotropy and Heterogeneity

The MR Egger intercept did not provide evidence for horizontal pleiotropy in analyses including multiple SNPs. Cochrane’s Q statistic did indicate heterogeneity in the analysis of the causal effect of worry on AN. However, the I^2^ statistic (and associated confidence intervals) did not. In the multiple SNP analysis of the causal effect of AN on the quantitative anxiety disorder phenotype, heterogeneity was indicated by Cochrane’s Q and I^2^. A leave one out sensitivity analysis did not suggest an overriding influence of any individual SNP, and in all multiple SNP analyses the confidence intervals of each SNP estimate overlapped. There was no marked improvement in heterogeneity with MR Egger estimates, relative to IVW estimates, in any of the multiple SNP analyses. Collectively there is no evidence to support bias caused by horizontal pleiotropy in IVW estimates of the study (for more detail see Supplementary Material, Tables 3 - 5 and Figures 8 and 9).

## Discussion

This study introduced MR to the study of AN, to investigate bidirectional effects of anxiety phenotypes and AN. The results of our MR analyses suggest that the genetic correlation identified between worry and AN is at least partly driven by worry exerting a causal influence on AN. In contrast there was no evidence to support a causal effect of AN on worry. There was also no evidence for causal effects between anxiety disorder pathology and AN, or of a genetic correlation between these phenotypes.

The finding that non-specific worry (i.e. worry that is not particularly directed towards eating and weight-gain) exerts a causal effect on AN risk is consistent with findings from previous cross-sectional (62-64) and longitudinal (65) observational studies. It has been suggested that worry inhibits emotional processing, and hinders problem solving (62), leading to a dependence on less adaptive coping mechanisms. Alternatively the focus on eating and weight (44), and even the neurobiological effects of dietary restriction (10, 66), may serve to alleviate worry in individuals who develop AN. Another possibility is that the process of worrying may put individuals at risk for a range of psychopathologies, with the content of worry determining the specific disorder that develops. Individuals with AN have elevated worry generally, but concern is particularly heightened in relation to eating, weight and shape (67, 68). Such may result when individuals prone to worrying direct their attention towards eating and weight, to drive the severe dietary restriction that is characteristic of AN.

While the precise mechanisms by which worry exerts its causal effects on AN require further investigation, our findings highlight the potential utility of addressing worry in eating disorder prevention. Existing interventions largely do not target non-specific forms of worry, and instead address disordered eating/weight-associated cognition. Two recent reviews highlight the efficacy of a number of existing interventions (particularly dissonance-based, cognitive-behavioural based, healthy weight programmes, media literacy programmes), in reducing disordered eating behaviour, and eating disorder symptoms, in individuals identified as at risk of eating disorders (69, 70). Future trials might explore whether the addition of components that reduce worry enhance the beneficial outcomes of these existing interventions. Worry may be targeted by a variety of adjunctive therapies (71). Mindfulness modules may be particularly useful additions to existing interventions given mindfulness practice discourages automatic and habitual patterns of thinking, including worry (71), and is reported to reduce body dissatisfaction (72). The benefits of reducing worry are likely to extend beyond eating disorder prevention, given the relevance of worry to both anxiety and depression (71).

Worry being a shared feature of both AN and anxiety disorders could explain the absence of causal association between AN and anxiety disorders observed in this study. Both anxiety and AN phenotypes may be underpinned by the common process of worry, with the presence of one signalling heightened risk for the other. Confounding of the anxiety disorder – AN association by a common factor would explain why the MR finding does not converge with previous observational studies (21, 30, 73). The latter report associations between anxiety disorders and AN but are subject to confounding, which is minimised in MR.

### Strengths and Limitations

A strength of the study is the use of MR, an approach that minimises risks of confounding and reverse causality, to robustly address questions of aetiology using secondary data. Sources of bias in MR are different from those affecting traditional observational epidemiology. The result of this is that where inferences from MR studies and those using other methods are consistent, as is the case for effects of worry on AN risk, we may be more confident that inferences are valid (74). This is particularly so when bias operates in different directions across studies, which might be expected here given bias in two-sample MR is typically towards the null (36).

A limitation of the MR approach is that it makes a number of assumptions that cannot be fully tested. The risk of confounding is reduced as compared to within studies of traditional epidemiological design, however it remains possible. We could not verify whether the genetic instruments were associated with plausible confounders of the exposure-outcome association in our sample, given the use of summary data (36). It is also impossible to determine whether instruments are associated with outcomes through pathways other than via the exposure of interest (75). To reduce the risk of incorrect inferences we completed a number of sensitivity analyses when multiple genetic instruments were available, with each sensitivity analysis robust to different MR assumptions. The causal effect of worry on AN was supported by all but the MR Egger estimate, which was very imprecise. Furthermore, the absence of evidence for pleiotropy, and the lack of improvement in heterogeneity in the MR Egger versus IVW model, suggests the IVW model provided a better fit to the data (57). Using estimates of R^2^ we confirmed that the majority of variants supported a direction of effect from worry to AN. Furthermore, MR estimates (IVW and sensitivity analyses) completed with this majority subsample of variants were consistent with the inference that worry increases risk of AN. We reduced the threshold for the strength of association between genetic variant and exposure to complete multiple-variant analyses of causal effects of anxiety disorders and AN, and subsequent sensitivity analyses. Findings of these analyses were consistent with those of the single-variant analyses. There was little evidence for heterogeneity across SNP estimates in all multiple-variant analyses, further supporting the absence of bias due to horizontal pleiotropy (76).

We used the largest GWAS for each phenotype of interest to date to maximise power (77), which could explain the discrepancy with a prior MR analysis that did not observe a causal effect of worry on AN (37). The anxiety disorder and AN GWAS sample sizes remained relatively small however, limiting power to detect a genetic correlation (50), as well as causal effects (77), between the two. This situation is likely to have been exacerbated by the anxiety disorder GWAS identifying variants associated with five anxiety disorders, introducing noise into the genetic signal (47). The primary determinant of power in a MR analysis is instrument strength, or variance in the exposure explained by the genetic instruments (41). Instrument strength in respect of the anxiety disorder and AN exposures is low, given few SNPs were robustly associated with these exposures, even when the threshold for association was reduced. This is likely to result from low statistical power of the GWASs, due to their sample size (78). Reducing the threshold for instrument identification further would have improved power (41). However the use of additional instruments increases the potential for pleiotropy (31), particularly when these instruments are weak. The use of weak instruments also introduces bias into the MR estimate due to confounding factors explaining greater variation in exposure and outcome compared to the instruments (42). In the case of two-sample MR this bias is in the direction of the null (77). Given the limitations surrounding power it is possible meaningful genetic associations between, and causal effects of, anxiety disorders and AN went undetected. Future studies should explore such further, using larger GWASs (with greater power to detect meaningful associations between instrumental SNPs and exposures) as these become available.

## Conclusion

The current study provides evidence for a causal influence of worry on AN. This finding is consistent with outcomes of previous observational studies, and may inform directions for future AN research and intervention. The low genetic signal in anxiety disorder and AN GWASs means we were not able to adequately assess the causal influence of these phenotypes. GWAS sample sizes are constantly growing, hopefully allowing for identification of increasingly robust genetic instruments for anxiety disorders and AN. This in turn will minimise bias and improve power, for rigorous assessment of causality that (with appropriate triangulation) can improve understanding and outcomes of AN.

## Acknowledgements

We are grateful to the participants who contributed to each of the GWASs considered in our study, and the research staff who collected and made available the data we used. The views expressed in this article are those of the authors and do not reflect official positions of our institutions or funders.

## Contributors

ECL and MRM conceived the study. ECL conducted the analysis and drafted the initial manuscript. HS guided all stages of the analysis. All authors assisted with interpretation of study outcomes, refining of manuscript drafts, and approved the final manuscript. ECL is the guarantor and attests that all listed authors meet authorship criteria and that no others meeting the criteria have been omitted.

## Ethics Approval

Participants gave informed consent for study participation and data sharing, as described in articles detailing original GWASs for each phenotype. Additional ethics approval was not required for this study.

## Data Sharing

All summary data used in the current study are publicly available. The AN and Anxiety Disorder GWAS summary statistics are available from the Psychiatric Genomics Consortium: https://www.med.unc.edu/pgc/results-and-downloads. Summary statistics in respect of the worry GWAS have been made available for download by the Complex Trait Genetics group, at the Center for Neurogenomics and Cognitive Research: https://ctg.cncr.nl/software/summary_statistics.

## Patient and public involvement

The current research used secondary data and was therefore not informed by patient and public involvement. We encourage future research directed by findings of the current study to seek patient and public guidance.

## Transparency

The lead author affirms that this manuscript is an honest, accurate, and transparent account of the study being reported; that no important aspects of the study have been omitted; and that any discrepancies from the study as planned have been explained.

